# Active control modulates memory-related activity during encoding and encoding–retrieval similarity

**DOI:** 10.64898/2026.08.23.746434

**Authors:** Zhuolei Ding, Shuge Yuan, Jincheng Xu, Shudong Zhang, Simon Hanslmayr, Xun Liu, Mingxia Zhang

## Abstract

Self-directed learning allows learners to actively control their learning experience and has been shown to enhance memory compared with matched yoked learning. However, it remains unclear when and how active control modulates memory-related neural activity during learning and retrieval. We recorded electroencephalography (EEG) while participants encoded objects under active and yoked learning conditions and again during a delayed recognition test approximately 24 h later. We examined event-related potential (ERP) activity across earlier processing windows, including pre-stimulus slow potentials and early N2 activity, and later processing windows, including P300, late slow-wave, and post-stimulus slow-potential activity. We also examined encoding–retrieval similarity (ERS) between neural patterns during encoding and retrieval. Behaviorally, active control improved delayed recognition, with the advantage selectively expressed in detailed recognition. In the ERP analyses, earlier processing windows showed memory-related effects, with pre-stimulus slow-potential and N2 activity differentiating subsequently remembered from forgotten items, but were not modulated by active control. By contrast, active control modulated later memory-related ERP activity, with remembered–forgotten differences expressed during late stimulus-related and immediate post-stimulus processing only in the active condition. ERS showed a similar active-control modulation: memory-related encoding–retrieval pattern similarity was evident under active learning, but not under yoked learning, with this effect involving relatively late encoding and retrieval windows. Together, these findings support a constructive-processing account, suggesting that memory formation under active control depends more strongly on rich, detailed encoding representations that can be reinstated during retrieval.

## 1 Introduction

Learning can be guided either externally or by the learner’s own control. Experimental paradigms that capture such control, commonly termed self-directed learning (SDL), typically contrast an active condition, in which learners control the order and timing of study items, with a yoked condition, in which participants study the same items in a sequence and timing matched to those generated by a previous participant in the active condition. Across a wide range of materials, populations, and developmental stages, these paradigms have consistently shown that active control enhances subsequent memory relative to the yoked condition (Ruggeri et al., 2025; Ding et al., 2025; Fantasia et al., 2020). Because yoked designs equate stimulus exposure, pacing, and visual history, this advantage cannot be attributed to differences in perceptual input or study time. Instead, it suggests that active control alters memory-related processes engaged during learning. However, the neural mechanisms underlying the mnemonic effects of active control remain poorly understood.

Two broad accounts have been proposed to explain why active control benefits memory (Markant et al., 2014, 2016). The first is a state-dependent optimization account, according to which active control improves memory by allowing learners to align information intake with moments of heightened readiness, arousal, or attention (Gruber & Otten, 2010; Markant et al., 2014; Otten et al., 2010; Tyng et al., 2017). On this view, active control should impact earlier memory-related activity, including preparatory activity before stimulus onset and early stages of stimulus processing. The second is a constructive-processing account, according to which active control changes the quality of encoding itself. Under this account, choosing what to inspect is thought to involve monitoring, evaluation and goal maintenance. These self-directed operations may promote more elaborative and integrative processing of the study item, such as evaluating whether the information has been sufficiently encoded, relating it to the ongoing learning context and actively constructing semantic meaning, thereby supporting the formation of richer episodic representations (Gureckis & Markant, 2012; Desender et al., 2021). On this view, the impact of active control should be expressed during later memory-related activity, including late stages of stimulus processing and immediate post-stimulus processing after stimulus offset.

Functional magnetic resonance imaging studies have implicated a hippocampal-prefrontal network supporting memory formation in SDL (Voss, Gonsalves, et al., 2011; Voss, Warren, et al., 2011). However, due to its limited temporal resolution, it remains unclear whether active control primarily alters earlier or later memory-related activity during learning. Electroencephalography (EEG), by contrast, provides the temporal resolution necessary to track when memory-related neural activity emerges during encoding. In particular, subsequent memory effects (SMEs), defined as differences in neural activity during encoding between items that are subsequently remembered and those that are forgotten (Kim, 2011; Mecklinger & Kamp, 2023; Paller & Wagner, 2002; Paller et al., 1987), provide an index of memory formation. By comparing SMEs between the active and yoked conditions across temporally distinct EEG components, we can test whether the mnemonic effects of active control are primarily expressed during earlier preparatory and stimulus-processing stages or during later stimulus-processing and immediate post-stimulus stages.

While temporally localized SMEs characterize when memory-related activity emerges, they do not directly capture how neural representations formed during encoding are reinstated during retrieval. To complement this process-level perspective, encoding–retrieval similarity (ERS) quantifies the correspondence between item-specific neural patterns at encoding and retrieval, providing an index of representational reinstatement in distributed neural activity patterns (Ritchey et al., 2013; Staresina et al., 2012). Prior intracranial EEG work has shown that volitional learning enhances item-specific ERS relative to passive learning, suggesting that active control can increase the reinstatement of encoding-related representations (Estefan et al., 2021). However, that study did not directly test whether active control modulates memory-related ERS, that is, whether the remembered–forgotten difference in ERS differs between active and passive learning. Moreover, this evidence was obtained from intracranial recordings in patients with epilepsy and focused primarily on hippocampal theta activity. It therefore remains unclear whether active control modulates memory-related representational reinstatement in healthy participants, as measured with non-invasive scalp EEG.

The present study used a well-established SDL paradigm with active and yoked conditions to examine how active control influences memory and, critically, whether it modulates memory-related neural processes in a manner more consistent with state-dependent optimization or constructive processing. The task comprised an encoding (learning) phase followed by a delayed recognition test (retrieval) phase, with EEG recorded during both phases. Here, “encoding phase” refers to the entire learning session rather than only the period of stimulus presentation or stimulus processing. For the ERP analyses, EEG activity during this phase was examined in windows before stimulus onset, during stimulus presentation, and immediately after stimulus offset.

To capture the temporal dynamics of these processes, we conducted univariate ERP analyses during encoding and multivariate representational similarity analysis (RSA) across encoding and retrieval to assess ERS. Based on the state-dependent optimization account, active control is expected to enhance earlier memory-related activity. Accordingly, in the ERP analyses, we focused on pre-stimulus slow potentials as an index of preparatory state (Gruber & Otten, 2010; Otten et al., 2010) and early N2 components as an index of early stimulus processing (Folstein & Van Petten, 2008; Tarbi et al., 2011). In contrast, the constructive-processing account predicts that active control primarily modulates later memory-related activity. To test this account, we examined the P300 and late slow-wave activity linked to stimulus evaluation and elaborative or integrative processing (Polich, 2007; Kamp et al., 2017; Mecklinger & Kamp, 2023), as well as post-stimulus slow potentials linked to post-stimulus registration and early consolidation (Ben-Yakov & Dudai, 2011; Ben-Yakov et al., 2013; Cohen et al., 2015; Yeung & Summerfield, 2012). We tested whether active control influences SMEs across these components. At the representational level, we further examined whether active control modulates memory-related ERS and when such effects emerge over time. Together, these analyses allowed us to test whether and when active control modulates temporally distinct memory-related neural activity during the encoding phase and memory-related encoding–retrieval correspondence.

## 2 Methods

### 2.1 Participants

Forty healthy right-handed volunteers participated in the study. One participant was excluded because of equipment malfunction. Eleven additional participants were excluded from the ERP analyses due to an insufficient number of artifact-free trials per condition (less than ten, see the Analyses section for details). The final sample for the ERP analyses therefore consisted of 28 participants (14 females; mean age = 22.43 years, SD = 2.64). For the RSA analyses, one additional participant was excluded due to incomplete data in the test phase, resulting in a final RSA sample of 27 participants (14 females; mean age = 22.37 years, SD = 2.68). All participants had normal or corrected-to-normal vision and reported no history of neurological or psychiatric disorders. All procedures were performed in accordance with the ethical standards of the Declaration of Helsinki. The study was approved by the Institutional Review Board of the Institute of Psychology, Chinese Academy of Sciences (approval number: H17023). Written informed consent was obtained from all participants before participation.

The sample size was determined based on prior behavioral effect sizes reported in the SDL paradigm, as well as sample sizes commonly adopted in recent EEG studies of episodic memory. Previous work has reported an effect size of Cohen’s d of 1.03 (Markant et al., 2014), for which an a priori power analysis (G*Power 3.1) indicated that a relatively small sample of 10 would be sufficient to achieve 80% power (α = .05, paired t-test, two-tailed). However, we additionally considered sample sizes commonly adopted in recent EEG studies of memory (Cheng et al., 2023; Wu et al., 2022) and therefore targeted a final sample of approximately 25 participants. To ensure this target could be met after data exclusion due to EEG artifacts (e.g., insufficient artifact-free trials per condition), we recruited 40 participants.

### 2.2 Materials

Stimuli comprised 456 object images drawn from prior studies (Ding et al., 2021; Xue et al., 2023). Eight images were reserved for practice trials and excluded from subsequent analyses. The remaining 448 images were divided into a study pool and an unstudied lure pool, each containing 224 images (112 living and 112 nonliving objects). The study pool was further subdivided into 14 sets, with seven sets assigned to the active condition and seven to the yoked condition during the encoding phase. The unstudied 224 images served as novel lures during the recognition test. The assignment of image sets to specific conditions of encoding phase and test phase (studied objects vs. unstudied lure) was fully counterbalanced across participants.

### 2.3 Procedure

The experiment was adapted from the SDL paradigm used in previous work (Ding et al., 2025, 2026; Ruggeri et al., 2019; Voss, Gonsalves, et al., 2011) and modified to be suitable for EEG recording. Participants completed an intentional encoding task followed by a delayed recognition test approximately 24 h later. Continuous scalp EEG was recorded during both the encoding and recognition sessions.

Prior to the main encoding phase, participants completed a brief familiarization phase consisting of one active block and one yoked block to practice the task requirements. The procedure was identical to that used in the main encoding phase (see below).

The main object encoding phase (see Figure 1A) consisted of 14 study blocks, with seven active blocks and seven yoked blocks alternating across the session (A-Y-A-Y…). Each block lasted 180 s. At the beginning of each block, 16 objects were displayed in a 4 x 4 grid for 5 s before being hidden by occluders. In the active condition, participants freely selected which object to inspect and whether to revisit previously viewed items. Selections were made via mouse click on the occluded locations. For each trial, selecting an object triggered a 0.5 s red frame around the chosen location, followed by a 1.5 s central fixation (pre-stimulus). The selected object was then presented centrally for 1.5 s (stimulus presentation), followed by a jittered blank inter-trial interval (ITI) of 2–3s before the next trial (post-stimulus). After an item was accessed, its occluder was replaced by a gray filler to indicate that it had already been learned. In the yoked condition, participants viewed the same sequence of item presentations as that generated by a previous participant in the active condition. On each trial, after a delay corresponding to the previous active participant’s response time on that trial, a red frame highlighted the to-be-revealed location, and participants clicked that location to remove the occluder. As in the active condition, the click triggered a 0.5 s red frame. The subsequent pre-stimulus fixation period, stimulus presentation, and post-stimulus jitter were identical to those in the active condition. After each active-yoked block pair, participants rested for at least 20 s and could extend the break if needed.

**Figure 1.**
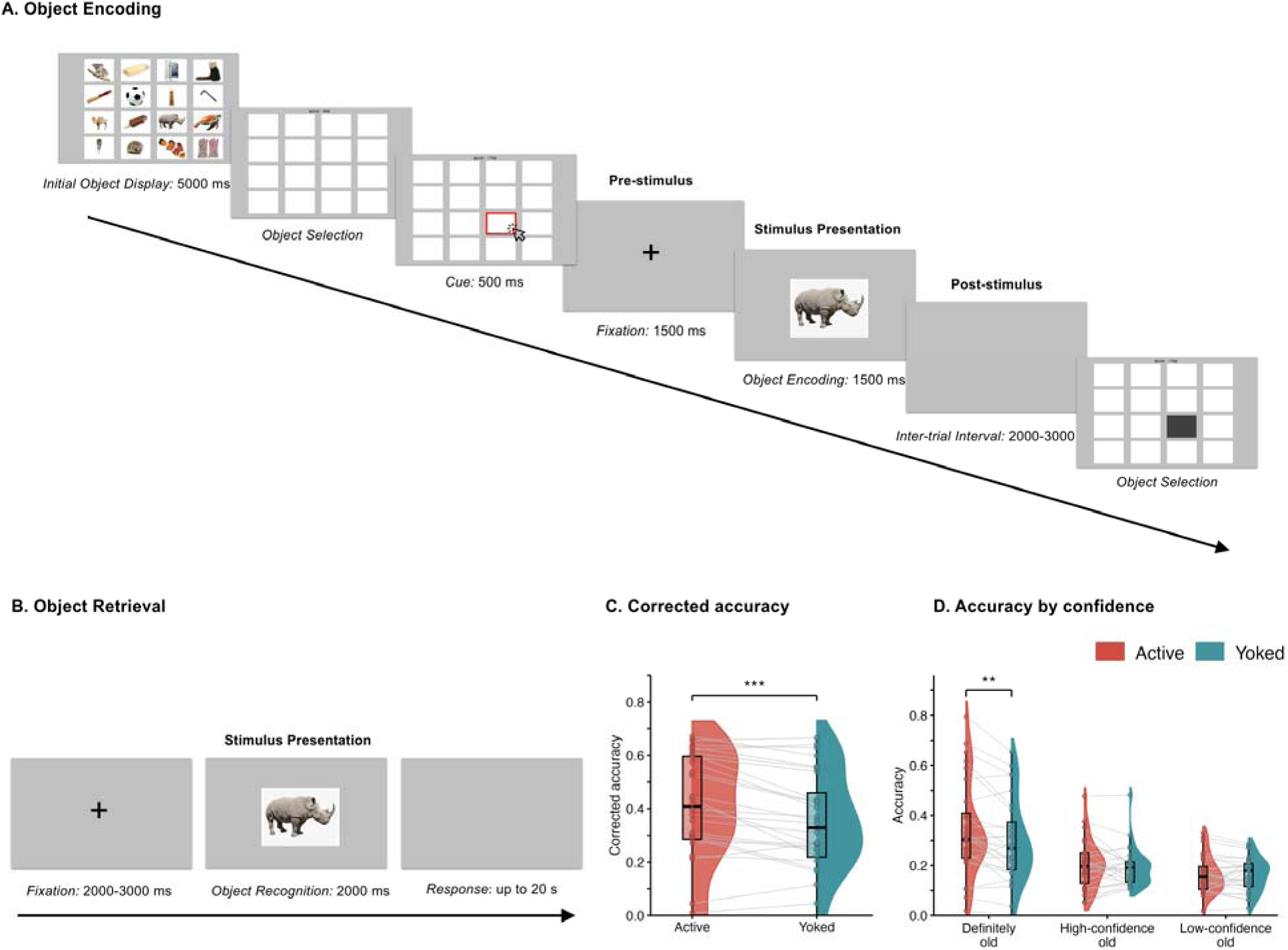
Experimental procedure and behavioral recognition performance in the active and yoked conditions. *Note.* **A.** Object encoding procedure. At the beginning of each block, all object images were displayed in a grid and then hidden by occluders. In the active condition, participants freely selected which object to inspect and whether to revisit previously viewed items, whereas in the yoked condition they followed the study sequence generated by a previous participant in the active condition. ERP analyses were time-locked to object onset and examined pre-stimulus, stimulus-presentation, and post-stimulus activity. **B.** Object retrieval procedure. Participants completed old/new recognition judgments with confidence for each object. ERS analyses quantified the similarity between item-specific neural patterns during stimulus presentation at encoding and retrieval. **C.** Corrected accuracy (hit rate minus false alarm) for studied objects in the active and yoked conditions. D. Accuracy by confidence in the active and yoked conditions. Dots represent individual participants, and grey lines connect observations from the same participant across the active and yoked conditions. Boxplots indicate the median and interquartile range, with whiskers extending to 1.5 times the interquartile range. Behavioral results are based on the ERP sample (*N* = 28). \*\**p*< .01, \*\*\**p* < .001.

Approximately 24 h later, participants returned for a recognition memory test (see Figure 1B). Eight practice trials were administered prior to the test to ensure that participants understood the task requirements. Each trial began with 2–3 s jittered fixation period. An object was then presented for 2 s, followed by a response window of up to 20 s. Participants rated their memory confidence on a 6-point recognition scale that ranged from definitely-old judgements to definitely-new judgements, with intermediate responses reflecting low confidence. The response options were: 1 = definitely-old with specific episodic details from the encoding episode, such as perceptual details of the object or contextual details including its grid location, presentation order, neighboring items, number of study repetitions, or encoding condition; 2 = high-confidence-old without retrieval of such specific details; 3 = low-confidence-old; 4 = low-confidence-new; 5 = high-confidence-new; and 6 = definitely-new. The recognition test consisted of randomized old and new trials presented across four test blocks separated by three 1-min breaks.

### 2.4 EEG Recording and Preprocessing

EEG data were recorded using a Brain Products system (Brain Products GmbH, Gilching, Germany) from 64 Ag/AgCl scalp electrodes positioned according to the extended 10–20 system during both the encoding (learning) phase and the delayed recognition (retrieval) phase. Participants were seated approximately 60 cm from a display screen in a sound-attenuated room. All channels were online referenced to FCz. Vertical electrooculogram (EOG) activity was recorded from an electrode placed below the right eye to monitor eye movements and blinks. Signals were sampled at 500 Hz, and electrode impedances were kept below 5 kΩ.

Offline preprocessing was conducted using custom MATLAB (The MathWorks Inc., 2013) and EEGLAB (Delorme & Makeig, 2004). The online reference at FCz was reinstated and included as a scalp channel. Continuous EEG data were re-referenced to the average of the left and right mastoids, consistent with prior ERP studies of subsequent memory effects (Gruber & Otten, 2010; Otten et al., 2010; Kamp et al., 2017), and band-pass filtered from 0.1 to 40 Hz. During the encoding phase, EEG epochs were created for stimulus-locked ERP averages, extending from -1500 ms to 2500 ms relative to stimulus onset. This extended time window allowed us to capture neural activity spanning pre-stimulus (−1500 to 0 ms), stimulus presentation (0 to 1500 ms), and post-stimulus stages (1500 to 2500 ms), with specific time windows and components of interest defined a priori (see EEG Data Analysis below). For baseline correction, we used the final 500 ms of the jittered blank ITI at the end of each trial. This baseline was chosen to avoid using the pre-stimulus fixation period as baseline, because that period was itself included in the analysis of pre-stimulus activity. During the retrieval phase, EEG epochs were extracted for stimulus-locked ERP averages, extending from −800 ms to 2000 ms relative to stimulus onset, with a baseline defined as the −500 ms interval preceding stimulus onset. To ensure high signal quality, multiple artifact rejection procedures were applied before averaging. First, epochs containing gross artifacts (e.g., excessive movement or amplifier saturation) were identified and removed. Second, independent component analysis (ICA) was performed to detect and remove artifacts associated with muscle, eye movements, and eye-blinks. Third, following ICA correction, trials were rejected if they contained peak-to-peak deflections exceeding ±100 µV.

### 2.5 EEG Data Analysis

#### ERP

ERP analyses were conducted on artifact-free epochs from the encoding phase, categorized according to Condition (Active vs. Yoked) and Memory (Remembered vs. Forgotten). Similar to previous studies (Gruber et al., 2013; Gruber & Otten, 2010), confidence ratings were taken into account to reduce the influence of guessing. Trials receiving low-confidence responses were excluded from the ERP analyses. Remembered trials were defined as studied items later judged as definitely-old or high-confidence-old, corresponding to ratings of 1 or 2. Forgotten trials were defined as studied items later judged as new. Participants with fewer than 10 artifact-free trials in any condition were excluded from further analyses (see Participants section for details). For the final sample for the ERP analyses, the mean number of artifact-free trials per participant was 72 (*SD* = 35.59) for Active Remembered, 30 (*SD* = 13.43) for Active Forgotten, 58 (*SD* = 27.20) for Yoked Remembered, and 35 (*SD* = 16.86) for Yoked Forgotten.

To test whether and when active control modulated memory-related activity, we selected five ERP components and time windows based on prior studies linking these signals to distinct stages of memory-related processing, as described in the Introduction. The pre-stimulus slow potential and N2 were selected as ERP components reflecting earlier processing stages. Specifically, earlier processing was analyzed over frontal electrodes (F1, Fz, and F2), including the pre-stimulus slow potential from −1000 to 0 ms (Gruber & Otten, 2010; Koen et al., 2018) and the N2 from 200 to 300 ms (Folstein & Van Petten, 2008; Kopp et al., 2020). The P300, late slow-wave, and post-stimulus slow potential were selected as ERP components reflecting later processing stages. Specifically, later stimulus- and post-stimulus-related activity were analyzed over central electrodes (C1, Cz, and C2), including the P300 from 300 to 600 ms (Polich, 2007), the late slow-wave from 700 to 1500 ms (Kamp et al., 2017), and the post-stimulus slow potential from 1500 to 2500 ms (Ben-Yakov & Dudai, 2011).

For each planned ERP component, mean amplitudes were submitted to 2 x 2 repeated-measures analyses of variance (ANOVAs) with factors Condition (Active, Yoked) and Memory (Remembered, Forgotten). Main effects of Condition and Memory, as well as their interaction, were evaluated within each time window. When significant interaction effects were observed, follow-up simple-effects analyses were conducted to characterize the direction of these effects.

#### ERS

We further examined ERS calculated as the similarity between neural patterns elicited by the same item at encoding and retrieval, providing an index of item-level pattern reinstatement over time (Justus et al., 2023; Kobelt et al., 2025; Liu et al., 2021; Popal et al., 2019). Analyses were restricted to time windows within 0–1500 ms following stimulus onset during both the encoding and retrieval phase. Following prior studies using spatiotemporal pattern similarity and encoding–retrieval similarity analyses (Lu et al., 2015; Pacheco Estefan et al., 2019, 2021; Kobelt et al., 2025), EEG data were z-transformed across electrodes and time points. The transformed data were segmented into overlapping 500 ms windows (Pacheco Estefan et al., 2019, 2021) with a step size of 20 ms (Kobelt et al., 2025). Each window was labeled by its center time (e.g., the 0–500 ms window is denoted 250 ms), and all encoding and retrieval times are reported as window centers. Within each window, voltage values across 250 time points and 62 electrodes were concatenated into a single vector, preserving spatiotemporal pattern information. For each trial, encoding vectors were correlated with the corresponding retrieval vectors using Spearman rank correlations. Correlation coefficients were Fisher z-transformed, yielding ERS matrices reflecting similarity across all combinations of encoding and retrieval time windows (encoding time × retrieval time). This temporally resolved approach allowed us to track the emergence of ERS over time.

To parallel the ERP analyses, ERS values were analyzed using planned contrasts corresponding to the main effect of Memory, the main effect of Condition, and their interaction at each encoding × retrieval time bin. The main effect of Memory was tested by contrasting remembered and forgotten trials collapsed across the active and yoked conditions: [(Active remembered + Yoked remembered) vs. (Active forgotten + Yoked forgotten)]. The main effect of Condition was tested by contrasting active and yoked trials collapsed across subsequent memory performance: [(Active remembered + Active forgotten) vs. (Yoked remembered + Yoked forgotten)]. The Condition × Memory interaction was tested by comparing the ERS-based subsequent memory effect between conditions: [(Active remembered − Active forgotten) vs. (Yoked remembered − Yoked forgotten)].

Statistical significance was assessed using a cluster-based permutation procedure (Kobelt et al., 2025; Maris & Oostenveld, 2007; Wu et al., 2022). For each planned contrast, the trial labels relevant to that contrast were shuffled within participants 1,000 times while preserving the original number of trials per condition. For each permutation, the contrast of interest was recomputed at each encoding × retrieval time bin. Time bins exceeding an uncorrected threshold of p < .05 were identified, and neighboring significant bins were grouped into clusters. The cluster-level statistic was defined as the number of significant time bins within each cluster, and the maximum cluster statistic from each permutation formed the null distribution. Observed clusters were considered significant if their statistic exceeded the 95th percentile of the permutation-based null distribution. For clusters that survived permutation correction, ERS values were averaged within each significant cluster and used to calculate the effect size, Cohen’s *d*. When a significant Condition × Memory interaction cluster was observed, mean ERS values were extracted separately for Active Remembered, Active Forgotten, Yoked Remembered, and Yoked Forgotten trials. Post-hoc paired-samples *t*-tests were then conducted to examine whether ERS differentiated remembered from forgotten trials within the active and yoked conditions, respectively, thereby characterizing how active control modulated the encoding–retrieval correspondence associated with successful memory.

### 2.6 Preregistration

No part of this study was preregistered.

### 2.7 Code availability

Custom MATLAB code used for EEG preprocessing, the encoding–retrieval similarity (RSA) analysis, the cluster-based permutation testing, and figure generation is available from the corresponding author upon reasonable request.

## 3 Results

### 3.1 Behavioral Performance

Behavioral analyses reported in the main text focused on the participants included in the ERP analyses (N = 28), ensuring direct correspondence between the behavioral and electrophysiological findings. Overall corrected recognition accuracy, calculated as hit rate minus false-alarm rate, was higher for actively selected than for yoked items (Active: *M* = 0.390, *SD* = 0.220; Yoked: *M* = 0.331, *SD* = 0.209; see Figure 1C), *t*(27) = 4.45, *p* < .001, *d* = 0.84. To examine whether the active-control advantage differed across confidence levels, we conducted a 2 × 3 repeated-measures ANOVA on recognition accuracy separately calculated for each old-judgement type, with Condition (Active, Yoked) and Judgement Type (definitely-old, high-confidence-old, low-confidence-old) as within-subject factors. This analysis revealed a significant main effect of Condition, F(1, 27) = 19.82, *p* < .001, *η²_p_* = .423, a significant main effect of Judgement Type, F(1.24, 33.59) = 9.99, *p* = .002, *η²_p_* = .270, and a significant Condition × Judgement Type interaction, F(1.66, 44.91) = 4.64, *p* = .020, *η²_p_* = .147. Bonferroni-corrected follow-up comparisons showed that accuracy for definitely-old judgements was higher for active than for yoked items (Active: *M* = 0.343, *SD* = 0.192; Yoked: *M* = 0.293, *SD* = 0.168), *t*(27) = 3.96, *p*_adj_ = .001, *d* = 0.75. In contrast, accuracy for high-confidence-old judgments did not differ between conditions (Active: *M* = 0.202, *SD* = 0.098; Yoked: *M* = 0.193, *SD* = 0.078), *t*(27) = 0.79, *p*_adj_ = 1.000, *d* = 0.15, nor did accuracy for low-confidence-old judgments (Active: *M* = 0.164, *SD* = 0.075; Yoked: *M* = 0.163, *SD* = 0.067), *t*(27) = 0.06, *p*_adj_ = 1.000, *d* = 0.01. Thus, the active-control advantage was most expressed in definitely-old recognition accuracy (see Figure 1D). Results for the full behavioral sample (N = 40) showed the same pattern (see Supplementary Materials).

### 3.2 ERP Results

#### Pre-stimulus slow potential (−1000 to 0 ms; F1, Fz, F2)

Pre-stimulus slow-potential amplitude showed a significant main effect of Memory (*F*(1, 27) = 11.82, *p* = .002, *η^2^_p_* = .31), such that subsequently remembered items were associated with more negative amplitudes than subsequently forgotten items (remembered: *M* = −2.78 μV, forgotten: *M* = −0.87 μV; see Figure 2A–D). Neither the main effect of Condition (*F*(1, 27) = 2.75, *p* = .109, *η^2^_p_* = .09), nor the Condition × Memory interaction (*F*(1, 27) = 0.32, *p* = .575, *η^2^_p_* = .01) reached significance.

**Figure 2.**
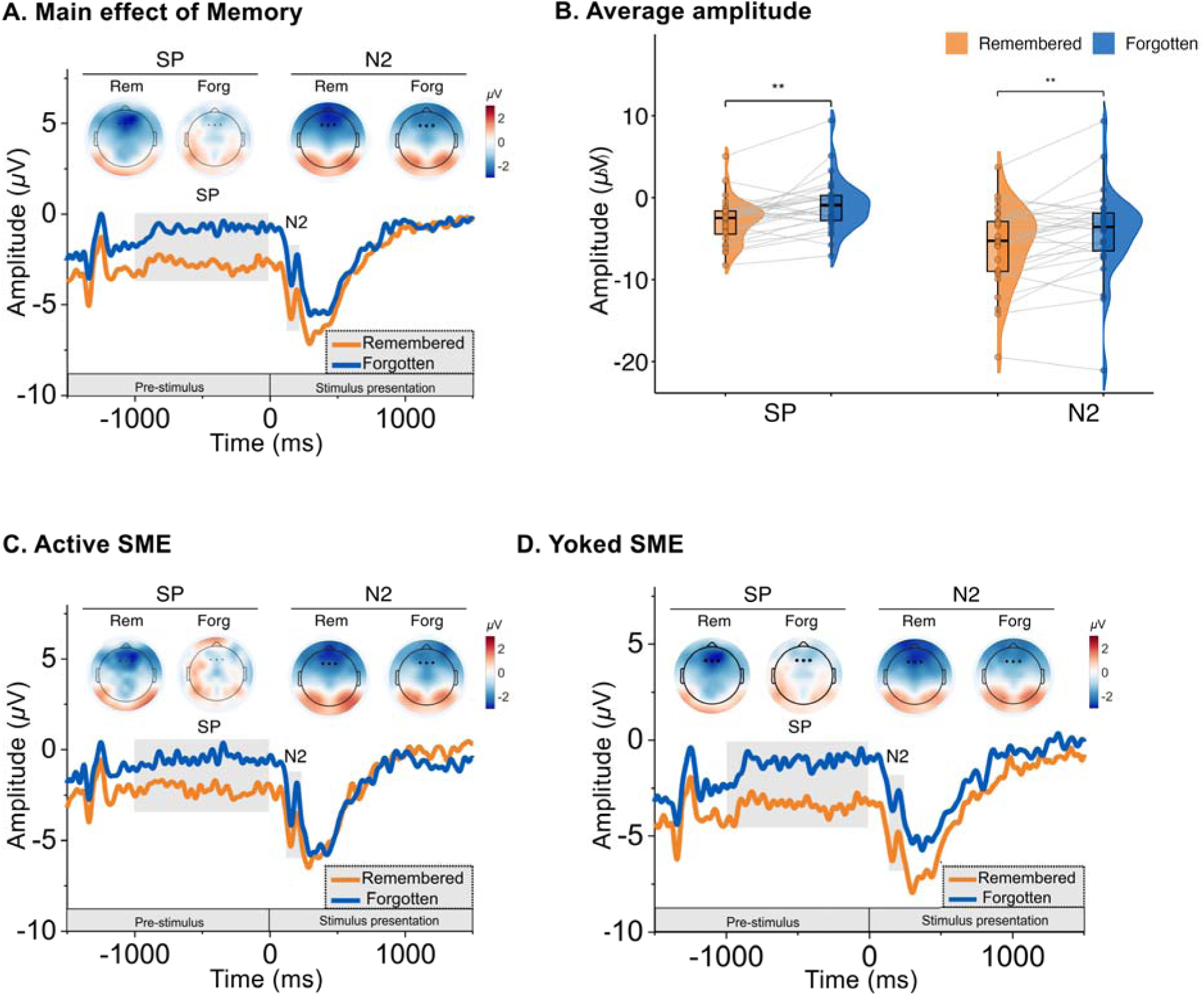
ERP: Subsequent memory effects during earlier processing windows. *Note*. **A.** Grand-average ERPs at frontal electrodes (F1, Fz, and F2) for subsequently remembered and forgotten objects, collapsed across conditions. Shaded areas indicate the pre-stimulus slow-potential (SP, −1000 to 0 ms) and N2 (200–300 ms) time windows and scalp maps show the corresponding voltage distributions. **B.** Mean amplitudes in the pre-stimulus slow-potential and N2 for subsequently remembered and forgotten objects, collapsed across conditions. Individual points represent participant-level mean amplitudes, and grey lines connect observations from the same participant across the subsequently remembered and forgotten objects. Boxplots indicate the median and interquartile range, with whiskers extending to 1.5 times the interquartile range. ** *p* < .01. **C.** and **D.**, Grand-average ERPs and scalp voltage distributions for subsequently remembered and forgotten objects in the active and yoked conditions, respectively.

#### N2 during stimulus presentation (200–300 ms; F1, Fz, F2)

The N2 component showed a significant main effect of Memory (*F*(1, 27) = 9.10, *p* = .006, *η^2^_p_* = .252), with more negative amplitudes for subsequently remembered items than forgotten items (remembered: *M* = −5.92 μV, forgotten: *M* = −4.18 μV; see Figure 2A–D). Neither the main effect of Condition (*F*(1, 27) = 0.75, *p* = .395, *η^2^_p_* = .027), nor the interaction between Condition and Memory was significant (*F*(1, 27) = 0.96, *p* = .337, *η^2^_p_* = .034).

#### P300 during stimulus presentation (300–600 ms; C1, Cz, C2)

The P300 showed a significant Condition × Memory interaction (*F*(1, 27) = 4.58, *p* = .042, *η^2^_p_* = .145, see Figure 3A), whereas neither the main effect of Condition (*F*(1, 27) = 0.41, *p* = .527, *η^2^_p_* = .015), nor that of Memory (*F*(1, 27) = 0.19, *p* = .664, *η^2^_p_* = .007), was significant. Follow-up simple-effects analyses suggested a tendency for remembered items to be associated with more positive amplitudes than forgotten items in the active condition (remembered: *M* = −2.60 μV, forgotten: *M* = −3.88 μV), although this difference did not reach significance (*t*(27) = 1.63, *p* = .116, *M*_diff_ = 1.27 μV, *SE*_diff_ = 0.78, 95% CI [−0.33, 2.88]). In contrast, no reliable difference was observed in the yoked condition (remembered: *M* = −3.90 μV, forgotten: *M* = −3.23 μV; *t*(27) = −0.79, *p* = .435, *M*_diff_ = −0.68 μV, *SE*_diff_ = 0.85, 95% CI [−2.43, 1.08]).Late slow-wave during stimulus presentation (700–1500 ms; C1, Cz, C2)

The late slow-wave showed a significant Condition × Memory interaction (*F*(1, 27) = 4.52, *p* = .043, *η^2^_p_* = .143), in the absence of significant main effects of Condition (*F*(1, 27) = 0.18, *p* = .675, *η^2^_p_* = .007), or Memory (*F*(1, 27) = 1.98, *p* = .171, *η^2^_p_* = .068). Simple-effects analyses showed that remembered items elicited more positive amplitudes than forgotten items in the active condition (remembered: *M* = 0.90 μV, forgotten: *M* = −0.74 μV; *t*(27) = 2.37, *p* = .025, *M*_diff_ = 1.63 μV, *SE*_diff_ = 0.69, 95% CI [0.22, 3.05]), whereas no difference was evident in the yoked condition (remembered: *M* = −0.30 μV, forgotten: *M* = −0.02 μV; *t*(27) = −0.45, *p* = .659, *M*_diff_ = −0.28 μV, *SE*_diff_ = 0.63, 95% CI [−1.57, 1.01]; see Figure 3A–D).

**Figure 3.**
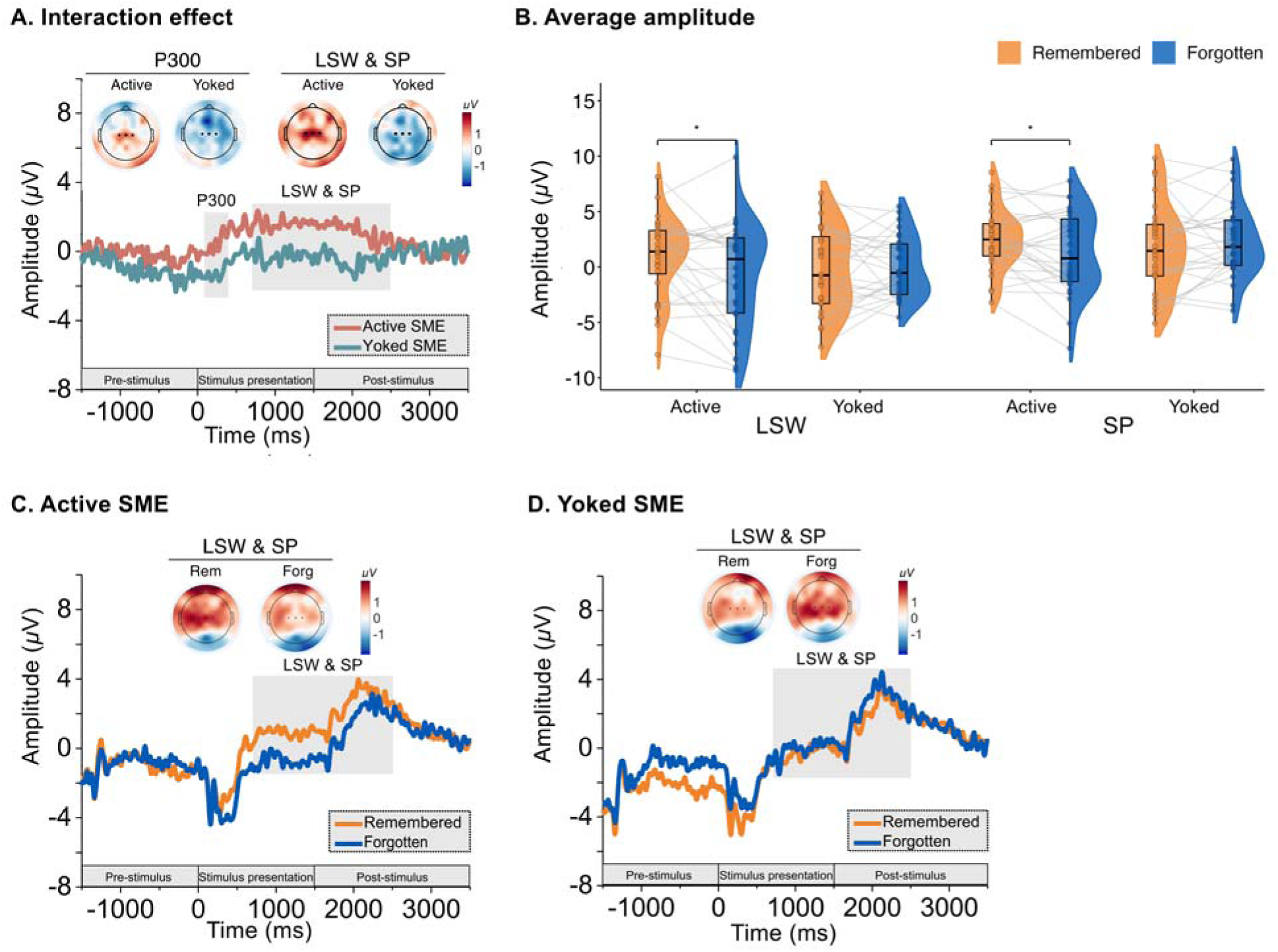
ERP: Condition-dependent subsequent memory effects during later processing windows. *Note*. **A.** Difference waveforms representing the SME at central electrodes (C1, Cz, and C2) in the active and yoked conditions. Shaded areas indicate the P300 (300–600 ms), late slow-wave (LSW, 700-1500 ms), and post-stimulus slow-potential (SP, 1500-2500 ms) time windows, and scalp maps show the corresponding SME voltage distributions. **B.** Mean amplitudes in the late slow-wave and post-stimulus slow-potential windows for subsequently remembered and forgotten objects in the active and yoked conditions. Individual points represent participant-level mean amplitudes, and grey lines connect observations from the same participant across subsequently remembered and forgotten objects in active and yoked condition. Boxplots indicate the median and interquartile range, with whiskers extending to 1.5 times the interquartile range. **C.** and **D.**, Grand-average ERPs for subsequently remembered and forgotten objects in the active and yoked conditions, respectively. * *p* < .05.

#### Post-stimulus slow potential (1500–2500 ms; C1, Cz, C2)

A significant Condition × Memory interaction was also found for post-stimulus slow-potential amplitude (*F*(1, 27) = 4.77, *p* = .038, *η^2^_p_* = .150), with no main effects of Condition (*F*(1, 27) = 0.04, *p* = .854, *η^2^_p_* = .001), or Memory (*F*(1, 27) = 0.80, *p* = .379, *η^2^_p_* = .029). Simple-effects analyses showed a significant difference between remembered and forgotten items in the active condition (remembered: *M* = 2.44 μV, forgotten: *M* = 1.23 μV; *t*(27) = 2.35, *p* = .026, *M*_diff_ = 1.21 μV, *SE*_diff_ = 0.52, 95% CI [0.15, 2.28]), whereas the corresponding comparison in the yoked condition was not significant (remembered: *M* = 1.66 μV, forgotten: *M* = 2.20 μV; *t*(27) = −0.92, *p* = .364, *M*_diff_ = −0.54 μV, *SE*_diff_ = 0.58, 95% CI [−1.74, 0.66]; see Figure 3A–D).

### 3.3 ERS Results

Cluster-based permutation tests revealed no significant clusters for either the main effect of Memory or the main effect of Condition (all *p*_cluster_ > .905). Importantly, a significant Condition × Memory interaction cluster emerged in the ERS matrix, spanning encoding windows from 250 to 950 ms and retrieval windows from 690 to 1250 ms (cluster size = 242 samples, *p*_cluster_ = .030, Cohen’s *d* = 0.55, 95% CI [0.16, 0.95]; see Figure 4A). This result indicates that memory-related ERS differed between the active and yoked conditions.

**Figure 4.**
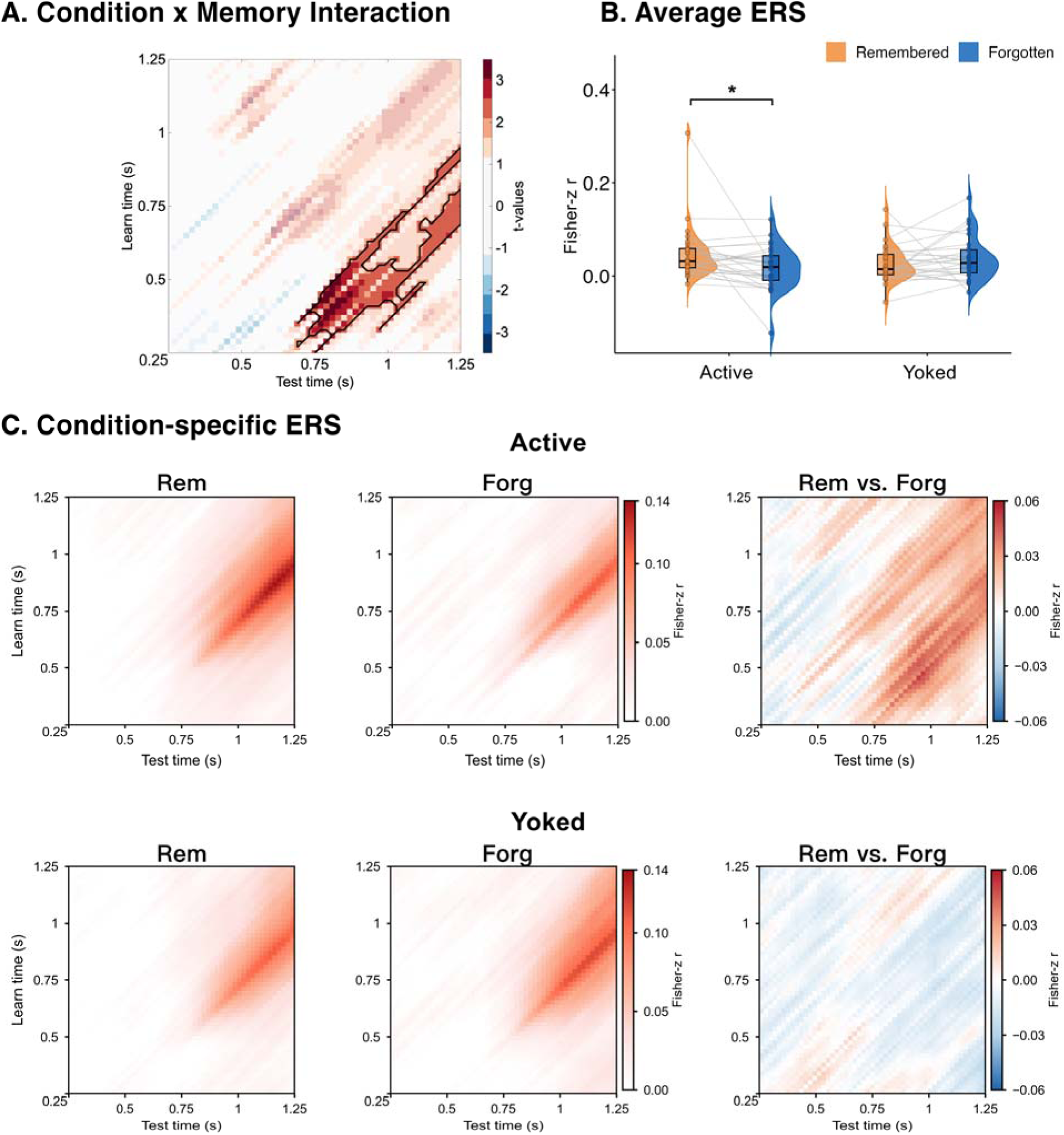
ERS: Condition-dependent memory-related effect. *Note.* ERS was computed as Fisher-z-transformed Spearman correlations between spatiotemporal EEG patterns at encoding and retrieval, using 500-ms sliding windows with a 20-ms step size. Both axes denote window-center times relative to stimulus onset (0.25–1.25 s); because each window is centered on its plotted time, the analyzed interval spans the full 0–1.5 s of stimulus presentation. **A.** Condition × Memory interaction. The black contour marks the significant cluster identified by cluster-based permutation testing. **B.** ERS averaged within the significant interaction cluster, shown separately for remembered and forgotten trials in the active and yoked conditions. Dots represent individual participants, and grey lines connect observations from the same participant across subsequently remembered and forgotten objects in active and yoked condition. Boxplots indicate the median and interquartile range, with whiskers extending to 1.5 times the interquartile range. **C.** Grand-average ERS for the four Condition × Memory cells, together with the corresponding remembered-minus-forgotten difference maps. \**p* < .05.

To characterize the direction of this interaction, mean ERS values were extracted from the significant cluster separately for Active Remembered, Active Forgotten, Yoked Remembered, and Yoked Forgotten trials. ERS was significantly higher for remembered than forgotten trials in the active condition (remembered: *M* = 0.05, *SD* = 0.06; forgotten: *M* = 0.02, *SD* = 0.05; *p* = .025; see Figure 4B). In contrast, ERS did not differ between remembered and forgotten trials in the yoked condition (remembered: *M* = 0.03, *SD* = 0.04; forgotten: *M* = 0.04, *SD* = 0.05; *p* = .328). Condition-specific ERS maps and the corresponding remembered–forgotten difference maps are shown in Figure 4C.

## 4 Discussion

The present study examined how active control influences memory and whether it modulates memory-related neural activity during encoding and memory-related encoding–retrieval similarity. Three main findings emerged. First, behaviorally, active control improved delayed recognition, and this advantage was selectively expressed in definitely-old responses. Second, in the ERP analyses, pre-stimulus slow potentials and early N2 activity differentiated subsequently remembered from forgotten items, but these earlier memory-related effects were not modulated by active control. In contrast, active control modulated later ERP memory-related effects: remembered items elicited more positive late slow-wave and post-stimulus slow-potential amplitudes than forgotten items in the active condition, whereas these remembered–forgotten differences were absent in the yoked condition. Third, in the ERS analyses, active control modulated memory-related ERS: ERS differentiated remembered from forgotten trials in the active condition, but not in the yoked condition. Together, these findings suggest that active control did not primarily alter earlier memory-related activity, including preparatory and early stimulus-related signals. Instead, its neural effects were expressed in later memory-related ERP activity, including late stimulus-related and immediate post-stimulus activity, as well as in memory-related ERS.

The frontal pre-stimulus slow potential and N2 both differentiated remembered from forgotten items, irrespective of whether learning was active or yoked. The frontal pre-stimulus slow potential is generally interpreted as an index of preparatory state, reflecting anticipatory fluctuations in attention and encoding readiness that precede an event (Weidemann & Kahana, 2021; Gruber & Otten, 2010; Otten et al., 2010). The frontal N2, in turn, has been associated with early attentional orienting and engagement to salient stimuli during initial stimulus processing (Folstein & Van Petten, 2008). The present earlier memory-related effects are consistent with the broader view that both pre-stimulus neural state (Guderian et al., 2009) and early stimulus-related processing (Mecklinger & Kamp, 2023) can contribute to subsequent memory.

Importantly, however, active control did not selectively strengthen these earlier memory-related effects. This finding is noteworthy because active control has often been proposed to enhance learning by increasing intrinsic motivation or the reward value, potentially by satisfying learners’ need for autonomy or agency (Ding et al., 2021; DuBrow et al., 2019; Leotti & Delgado, 2011; Murty et al., 2015). Previous work on reward and memory has shown that monetary reward anticipation can increase pre-stimulus neural activity and modulate pre-stimulus SMEs (Adcock et al., 2006; Gruber & Otten, 2010; Poh et al., 2022; Shohamy & Adcock, 2010). Evidence from other domains, such as decision making further suggests that N2 activity can be sensitive to reward and motivational significance (Novak & Foti, 2015). In the present study, however, active control did not modulate either the pre-stimulus slow-potential or the N2 subsequent memory effect. Thus, although active control may share motivational properties with reward, its influence on memory does not appear to be primarily expressed as a reward-like enhancement of preparatory or early stimulus-related memory processes.

By contrast, active control modulated later ERP memory-related activity, including late stimulus-related and immediate post-stimulus activity. In the P300 window, active control already modulated memory-related activity, although the remembered–forgotten difference within the active condition was only marginal. The effect became more robust in later time windows: in both the late slow-wave and post-stimulus slow-potential periods, subsequently remembered items elicited more positive amplitudes than forgotten items in the active condition, whereas no remembered–forgotten difference was observed in the yoked condition. Thus, the ERP effects of active control were expressed most consistently in time windows associated with late stimulus processing and immediate post-stimulus processing, rather than in pre-stimulus preparatory or early stimulus-related activity.

The functional interpretation of these later ERP effects can be grounded in prior work on positive-going ERP subsequent memory effects. The P300 has been linked to the allocation of attentional resources and the updating of stimulus-context representations during stimulus evaluation (Polich, 2007; Mecklinger & Kamp, 2023). Later positive slow-wave activity is often associated with sustained elaborative, organizational, or integrative encoding operations that support later remembering (Mecklinger & Kamp, 2023; Kamp et al., 2017). The post-stimulus slow potential further suggests that memory-related processing may continue after stimulus offset, potentially involving the registration, evaluation and early stabilization of newly encoded events (Ben-Yakov & Dudai, 2011; Ben-Yakov et al., 2013; Cohen et al., 2015; Mecklinger & Kamp, 2023). In the current study, active learning may have involved greater monitoring, evaluation and goal maintenance, allowing the selected item to become embedded within a self-generated learning context. This, in turn, may have promoted more elaborative processing, such as evaluating the item’s encoding status, relating it to the ongoing learning episode, integrating it with a self-generated study sequence, and constructing semantic or contextual links, all of which may contribute to memory formation (Gureckis & Markant, 2012; Markant et al., 2016; Yeung & Summerfield, 2012; Kamp et al., 2017). These findings complement prior behavioral and fMRI work showing that choice and volitional exploration enhance memory and recruit hippocampal–prefrontal circuitry (Markant et al., 2014, 2016; Murty et al., 2015; Ding et al., 2021, 2024; Voss, Gonsalves, et al., 2011), and provide temporal evidence consistent with a constructive-processing account: self-directed choice may promote memory by engaging later elaborative and integrative encoding.

Furthermore, the ERS analysis showed that active control modulated memory-related encoding–retrieval pattern similarity. Specifically, ERS differentiated subsequently remembered from forgotten items in the active condition, whereas no comparable remembered–forgotten ERS difference was observed in the yoked condition. This finding suggests that self-directed learning not only modulated memory-related ERP activity during encoding, but also influenced the extent to which neural patterns established during encoding were recovered during later retrieval. Notably, this ERS effect involved both encoding- and retrieval-side time windows. On the encoding side, the effect was centered between 250 to 950 ms after stimulus onset, a period that overlaps with the P300 window and a portion of the late slow-wave window. On the retrieval side, the effect was centered between 690 and 1250 ms after stimulus onset, a relatively late time window that is more consistent with the reinstatement of encoding-related representations than with early sensory analysis (Staresina & Wimber, 2019).

How might active control give rise to this memory-related ERS? Building on the late slow-wave and post-stimulus ERP findings, one possibility is that successful remembering under self-directed learning depends more on the formation of encoding representations that are distinctive, well organized, and tightly linked to the learner’s self-generated study context (Markant et al., 2016; Gureckis & Markant, 2012; Yeung & Summerfield, 2012). Such representations may be more likely to be re-expressed when the item is later encountered, allowing ERS to differentiate remembered from forgotten items in the active condition. This interpretation is consistent with evidence that successful remembering depends not only on the strength of encoding, but also on the overlap between neural patterns present during encoding and those reinstated during retrieval (Ritchey et al., 2013; Gordon et al., 2014; Hill et al., 2021). The behavioral findings fit the same logic: the active-control benefit was expressed specifically in detailed old recognition, that is, “definitely old” responses accompanied by retrieval of specific episodic details from the encoding episode, rather than in high-confidence old responses without such details or low-confidence old responses. This pattern suggests that self-directed encoding may have supported richer, more detailed representations rather than merely increasing a nonspecific sense of familiarity. Together, the ERP, ERS, and behavioral findings converge on a constructive-processing account in which active control supports the formation of richer encoding representations that are more likely to be reinstated during detailed retrieval.

## 5 Conclusion and Limitations

The present study shows that active control during learning enhanced memory and shaped both memory-related ERP activity during encoding and memory-related encoding–retrieval pattern similarity. Earlier memory-related effects, including pre-stimulus slow-potential and early N2 effects, differentiated subsequently remembered from forgotten items regardless of whether learners controlled the study sequence. In contrast, active control modulated later memory-related ERP activity, with remembered–forgotten differences expressed during late stimulus-related and immediate post-stimulus processing in the active condition only. The ERS findings further support this interpretation: memory-related encoding–retrieval pattern similarity differentiated remembered from forgotten items only under active learning. Together with the selective behavioral effect on detailed recognition, these findings converge on a constructive-processing account in which active control supports the formation of richer encoding representations that are more likely to be reinstated during retrieval.

Two limitations should be noted. First, active control itself may involve multiple self-directed processes, such as strategic monitoring, study-sequence planning, revisiting, and mnemonic integration (Markant et al., 2014, 2016; Ding et al., 2026). These processes may contribute to the later memory-related ERP activity and memory-related ERS observed here in different ways, but the present study cannot determine their respective contributions. Future work could experimentally isolate different components of active control to test how each process, alone or in combination with others, influences memory. Second, although the behavioral results showed that the active-control benefit was selectively expressed in detailed old recognition, the ERP and ERS analyses were designed to compare remembered and forgotten trials and did not further subdivide remembered trials by confidence or retrieval of specific episodic details. Although the present study included a relatively large number of trials and participants for a memory EEG study (224 trials per participant; N = 40), the number of artifact-free trials remained insufficient for stable neural comparisons between detailed and non-detailed old recognition. Future studies with larger samples, more trials, or multi-session designs could directly compare these two forms of remembering.

## Author contributions

**Z.D.**: Software, Formal analysis, Writing – original draft, Writing – review & editing, Visualization. **S.Y.**: Formal analysis, Writing – original draft, Writing – review & editing, Visualization. **J.X.**: Investigation, Data curation. **S.Z.**: Supervision. **S.H.**: Supervision, Writing – review & editing. **X.L.**: Supervision. **M.Z.**: Conceptualization, Writing – review & editing, Supervision, Funding acquisition.

## Data availability

The de-identified participant-level data supporting the findings of this study (trial-averaged ERP amplitudes and participant-level encoding–retrieval similarity matrices) are available on the Open Science Framework at https://osf.io/2ne3h/overview?view_only=980eda1f31414c5386f21151d2961028. Because the raw EEG recordings constitute potentially identifiable human data collected under participant consent that does not permit fully open release, raw EEG data are available from the corresponding author under a data use agreement for non-commercial research purposes, subject to approval by Institute of Psychology, Chinese Academy of Sciences IRB.

## Preregistration

No part of this study was preregistered.

## Ethics approval and consent to participate

This study was performed in accordance with the ethical standards of the Declaration of Helsinki. The study protocol was reviewed and approved by the Institutional Review Board of the Institute of Psychology, Chinese Academy of Sciences (approval number: [H17023]). Written informed consent to participate and to the publication of anonymised data was obtained from all participants prior to the start of the experiment.

## Competing interests

The authors declare no competing interests.

## Funding information

M.Z. discloses support for the research of this work from the National Natural Science Foundation of China [62177045]. Z.D., S.Y., J.X., S.Z., S.H. and X.L. declare no relevant funding.

## Notes

### Competing Interest Statement

The authors have declared no competing interest.

