## Appendix for "Active control modulates memory-related activity during encoding and encoding–retrieval similarity"

**Supplementary Materials**

**Behavioral analyses of the full behavioral sample (N = 40)**

Corrected recognition (hits minus false alarms) was higher for active than yoked items (Active: *M* = 0.408, *SD* = 0.224; Yoked: *M* = 0.355, *SD* = 0.210), *t*(39) = 4.57, *p* < .001, *d* = 0.72. To examine whether this active-control advantage differed across old-response types, we conducted a 2 × 3 repeated-measures ANOVA on old-item response proportions, with Condition (Active, Yoked) and Judgement Type (definitely-old, high-confidence-old, low-confidence-old; ratings 1, 2, and 3, respectively) as within-subject factors. Greenhouse-Geisser correction was applied to effects involving Judgement Type. This analysis yielded a significant main effect of Condition, *F*(1, 39) = 20.91, *p* < .001, η²p = .349, a significant main effect of Judgement Type, *F*(1.20, 46.82) = 12.85, *p* < .001, η²p = .248, and a significant Condition × Judgement Type interaction, *F*(1.72, 67.20) = 5.09, *p* = .012, η²p = .115. Bonferroni-corrected follow-up comparisons showed that definitely-old responses were more frequent for active than yoked items (Active: *M* = 0.343, *SD* = 0.198; Yoked: *M* = 0.301, *SD* = 0.182), *t*(39) = 3.98, *p*_adj_ < .001, *d* = 0.63. In contrast, high-confidence-old responses did not differ between conditions (Active: *M* = 0.208, *SD* = 0.100; Yoked: *M* = 0.196, *SD* = 0.089), *t*(39) = 1.34, *p*_adj_ = .564, *d* = 0.21, nor did low-confidence-old responses (Active: *M* = 0.166, *SD* = 0.072; Yoked: *M* = 0.167, *SD* = 0.072), *t*(39) = -0.18, *p*_adj_ = 1.000, *d* = -0.03.
